# *Trans*-Translation inhibitors and copper synergize for enhanced antibiotic activity

**DOI:** 10.1101/2024.11.14.623546

**Authors:** Melissa Vázquez-Hernández, Huriye Deniz Uzun, Mona Otte, Lea Schipp, Christoph H. R. Senges, Sascha Heinrich, John N. Alumasa, Jennifer Stepien, Fatih Demirbas, Paolo Cignoni, Katrin Marcus-Alic, Kristina Tschulik, Ute Krämer, Thomas Günther Pomorski, Kenneth C. Keiler, Julia E. Bandow

## Abstract

*Trans*-Translation is the most effective ribosome rescue mechanism and a compelling target for new antimicrobial agents. A recent proteomic study revealed similarities between the responses of *Bacillus subtilis* to the inhibitors small-molecule inhibitors oxadiazole KKL-40 and tetrazole KKL-55 and divalent cation ionophores, indicating the disturbance of metal homeostasis as potential secondary mechanism of action. Here, we report increased copper levels in KKL-40 and KKL-55-treated *B. subtilis*. Both inhibitors form copper complexes that enter large unilamellar vesicles. Copper supplementation enhanced the antibacterial activity against *B. subtilis* by simultaneously increasing inhibitor and copper uptake. The co-treatment of *B. subtilis* with *trans*-translation inhibitors and copper concentrations normally benign for *trans*-translation-competent cells, caused an immediate stalling of growth and translation, as observed at higher KKL-40 and KKL-55 concentrations without copper supplementation. Proteome analysis showed that during translation stalling cells were unable to mount an effective copper toxicity response. Taken together, the synergetic uptake of KKL-40 and KKL-55 with copper leads to a quick-onset translation stalling, preventing *B. subtilis* from counteracting the toxic effects of rapid copper influx.

**Significance statement:** The challenge of antimicrobial resistance is growing, necessitating an exploration of novel antibiotic targets. Among these, *trans*-translation has attracted considerable attention due to its ubiquitous presence in bacteria as well as its role in virulence and pathogenesis. Several inhibitors of *trans*-translation have been identified in a target-based screening using a whole-cell assay. However, recent proteomic profiling studies suggested that the tested *trans*-translation inhibitors might have an additional mode of action. In this work, we shed light on their previously undiscovered copper ionophore activity and explore the consequences of co-treating *B. subtilis* with KKL-40 or KKL-55 and CuCl_2_. This co-treatment results in a rapid antibiotic influx, and, consequently to the stalling of ribosomes, translation, and bacterial growth. Simultaneously, massive amounts of copper accumulate in the cells, the toxic effects of which require a copper stress response to mitigate. However, such a response is averted by the stalled translation. Dual mechanism antibacterial agents are attractive because they are typically associated with slow emergence of resistance. A deep understanding of the complex interplay of KKL-40 and KKL-55 with metal ions will help to fully exploit *trans*-translation as an antibacterial target and to develop KKL-40 and KKL-55-based antibiotics into novel treatments for bacterial infections.

## Introduction

A study published in early 2022 assessed the impact of bacterial antimicrobial resistance (AMR) (1). It revealed that nearly 5 million deaths in 2019 were associated with AMR, with over 1 million directly attributable to resistance. Approximately 70% of those deaths were caused by pathogens resistant to fluoroquinolones and ß-lactams, antibiotics in clinical use for decades. To address the AMR challenge, the search for new antimicrobials with novel targets and mechanisms of action is becoming a priority.

*Trans*-Translation, a mechanism for ribosome rescue in bacteria, has been identified as a promising target (2). Previous studies found that *trans*-translation is needed for viability or virulence in all pathogens tested (3–5). *Trans*-Translation depends on a ternary complex composed of tmRNA, SmpB, and EF-Tu·GTP that recognizes and releases stalled ribosomes from the mRNA while targeting both the trapped aberrant nascent peptide and the mRNA for degradation (6). Although alternative ribosome rescue systems are known in bacteria, such as ArfA and ArfB in *Escherichia coli*, ArfT in *Francisella tularensis*, and BrfA and a ribosome-associated quality control factor (RqcH) in *Bacillus subtilis*, *trans*-translation is the primary and most effective ribosome rescue system (6–11). In 2013, oxadiazole and tetrazole-based compounds were identified as broad-spectrum antibiotics capable of inhibiting *trans*-translation (12). The identified compounds showed antimicrobial activity against *B. subtilis, Bacillus anthracis*, *Staphylococcus aureus*, *Mycobacterium smegmatis*, *Mycobacterium tuberculosis*, *E. coli*, *F. tularensis, Legionella pneumophila*, and *Neisseria gonorrhoeae* (12–19). The molecular targets of oxadiazoles and tetrazoles differ: the oxadiazole KKL-35 was shown to target a unique site near the peptidyl-transferase center of the ribosome (19), while the tetrazole KKL-55 binds EF-Tu, preventing it from binding tmRNA (20).

These molecules inhibited *trans*-translation *in vitro* and *in vivo*, but the minimal inhibitory concentrations (MICs) were unchanged against *E. coli* strains deleted for *ssrA* (and therefore unable to perform *trans*-translation), suggesting a second mechanism of action (12). Similarly, KKL-35 inhibited growth of the *trans*-translation-deficient Δ*ssrA* and Δ*smpB* mutants of *L. pneumophila* (21), which would not be expected if *trans*-translation were the only target. Furthermore, a *B. subtilis* proteomic response library was used to compare the responses elicited by the oxadiazoles KKL-35 and KKL-40 and the tetrazole KKL-55 with those evoked by other antimicrobial agents. The proteomic responses to treatment with KKL-40 and KKL-55 were similar to those induced by the ionophores calcimycin, 4-Br-calcimycin, and ionomycin, which transport calcium, copper, iron, manganese, and/or zinc across biological membranes (13, 22).

In the present study, we used the gram-positive model organism *B. subtilis* to investigate the role of metals in the antibacterial mechanisms of KKL-40 and KKL-55 (see Fig. 1a for chemical structures). We assessed the impact of KKL-40 and KKL-55 on metal homeostasis using inductively-coupled plasma optical emission spectrometry (ICP-OES). Furthermore, we analyzed the capability of KKL-40 and KKL-55 to form metal complexes and mediate metal transport *in vitro*. Following the observation of synergetic transport, we evaluated the impact of *trans*-translation inhibitors and CuCl_2_ co-treatment on antibiotic efficacy, KKL and copper accumulation, and the bacterial response at the proteome level. We conclude that the rapid influx of high amounts of KKL leads to translation arrest, making it impossible for *B. subtilis* to mitigate the toxic effects of inadequate ribosome rescue capacity and of rapid copper influx.

**Figure 1.**
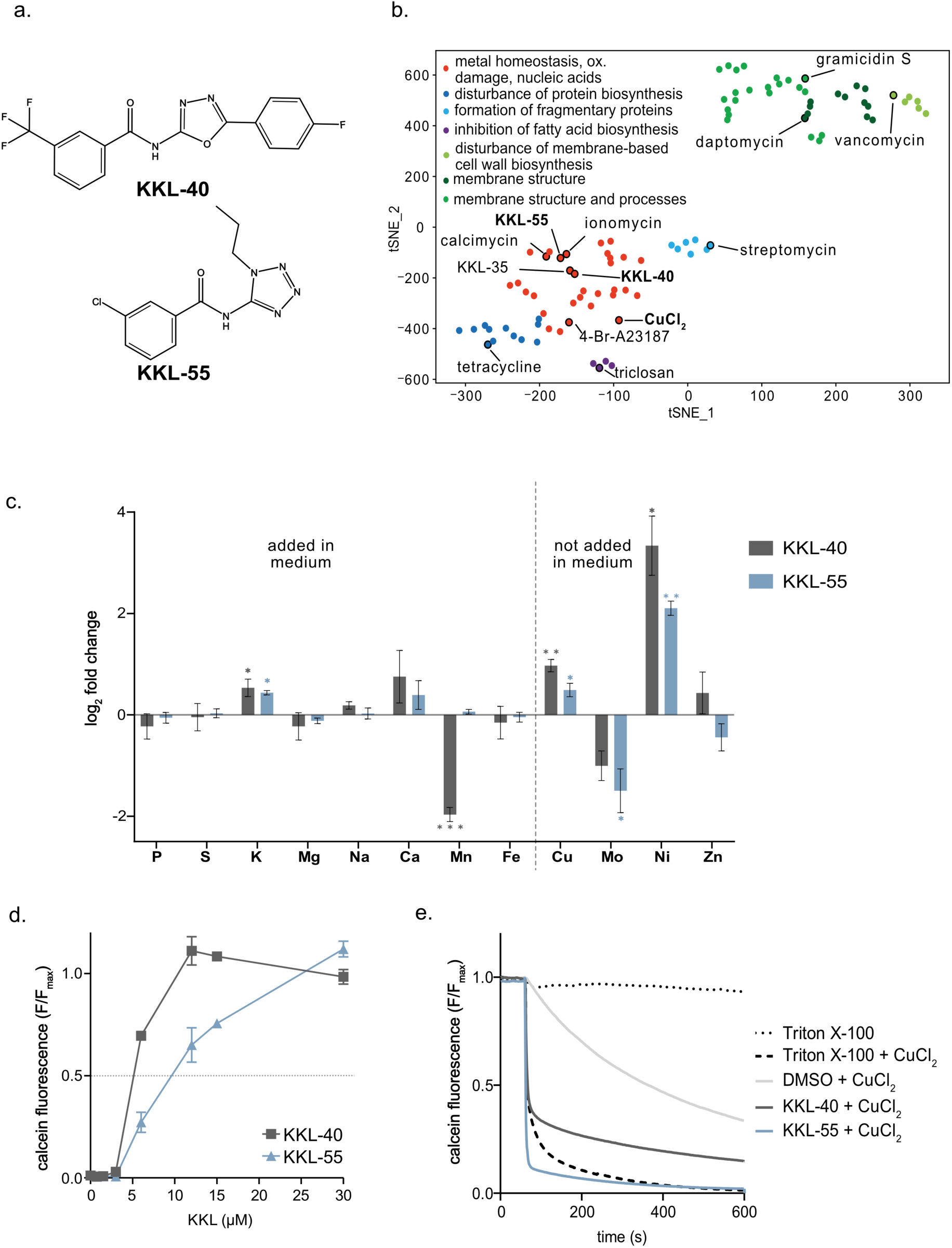
KKL-40 and KKL-55 elicit a proteomic response similar to ionophores, complex copper, and mediate copper transport into unilamellar vesicles. (a) Chemical structures of KKL-40 and KKL-55. (b) t-Distributed stochastic neighbor embedding of proteomic responses of *B. subtilis* 168 to different antibiotics, including KKL-40 and KKL-55 (13). Each dot represents a proteomic response of *B. subtilis* to an antibacterial agent. Compounds investigated in detail in this study are highlighted in bold. Proteomic data were sourced from previous publications (13, 22), with a fully annotated figure available in Supplementary File 1 Supplementary Fig. S1. (c) Changes in cellular element concentrations after 15 min of treatment with subinhibitory concentrations of 0.8 µM KKL-40 (dark grey) or 7.2 µM KKL-55 (blue) compared to untreated cells, determined by ICP-OES. Data represent the log_2_-transformed average fold-changes and standard deviations (SD) of three biological replicates. Statistical analyses were performed with Student’s *t*-test against the untreated control (^✱^ p ≤ 0.1; ^✱✱^ p ≤ 0.01; ^✱✱✱^ p ≤ 0.001). (d) The complex formation of KKL-40 and KKL-55 with Cu(II) was evaluated using calcein, a fluorescent dye quenched by Cu(II). Calcein (3 µM) was quenched by CuCl_2_ (3 µM). Inhibitors were added to reach final concentrations of 0.75, 1.5, 3, 6, 12, 15, or 30 µM. Fluorescence of calcein in the absence of CuCl_2_ was set to 1 (F_max_) and in the presence of 3 µM CuCl_2_ to 0. HEPES (50 mM, pH 7.4) was used as buffer. Data represent the means and SDs of three technical replicates. (e) *In vitro* copper transport into calcein-filled large unilamellar vesicles (LUV). LUVs were preincubated with 1 µM KKL-40 or KKL-55 for 5 min at 4°C before measuring fluorescence at 10°C. CuCl_2_ (3 µM) was added after 60 s, indicated by the arrow. Fluorescence quenching indicates copper influx. As controls, calcein-filled LUVs were also incubated with Triton X-100 at lytic concentration (0.2% v/v) in the absence or presence of CuCl_2_. Fluorescence was measured at λ_ex_ = 480 nm (λ_em_ = 520 nm) at 10°C. Representative data, normalized to the maximal fluorescence (F_max_) are shown here; additional replicates are shown in Supplementary File 1 Supplementary Fig. S5.

## Results and Discussion

### 1. KKL-40 and KKL-55 disturb metal homeostasis in *B. subtilis*

As part of a large comparative proteome analysis we recently reported proteomic responses of *B. subtilis* to the *trans*-translation inhibitors KKL-35 and KKL-40 (both oxadiazoles) and the structurally unrelated tetrazole KKL-55 (13). Rather unexpectedly, these showed similarities to the proteomic responses observed for the ionophores calcimycin and ionomycin. In contrast to target-centric assays, the proteomic responses mirror the global impact of a compound on bacterial physiology and may thus reveal secondary mechanisms or target areas (23, 24). Our expanding reference library of proteomic responses of *B. subtilis* (13, 25), together with a similarity matrix-based computational approach (26), allows us to identify commonalities between new antibiotics and previously tested ones (13). Following up on the ionophore-like proteome response profiles, in the present study we applied t-distributed stochastic neighbor embedding (t-SNE) (27) to the similarity matrix that was updated to include the response to the copper ionophore 4-Br-A23187 (22) (Fig. 1b, Supplementary File 1 Supplementary Fig. S1). The proximity of KKL-40 and KKL-55 to the carrier ionophores calcimycin and ionomycin in the t-SNE plot indicates a high similarity of the proteomic responses, reflecting the upregulation of a significantly overlapping set of proteins.

Ionophores mediate the transport of ions across membranes, thereby affecting metal homeostasis and/or the proton gradient of bacterial cells. To evaluate the ion homeostasis in *B. subtilis* after treatment with KKL-40 or KKL-55, concentrations of the elements P, S, K, Mg, Na, Ca, Mn, Fe, Cu, Mo, Ni, and Zn in the soluble fraction of cell lysate (containing the cytosolic contents and membrane fraction) were determined by ICP-OES after a 15-min treatment during early log-phase (Fig. 1c, Supplementary File 2). Cells were grown in the same chemically-defined medium (Belitzki minimal medium; BMM) used for the previously reported and the current proteome analyses, and treated with the concentrations of KKL-40 or KKL-55 that reduced the growth rate by ∼50% (0.8 µM (0.3 µg/ml) KKL-40, or 7.2 µM (2 µg/ml) KKL-55). Copper, molybdenum, nickel, and zinc are not actively added during the manufacture of the medium (BMM), so that the detected levels of these elements are due to impurities from other ingredients or materials. For most elements, concentrations remained unchanged (Fig. 1c, Supplementary File 2). However, after treatment with KKL-40 and KKL-55, copper and nickel levels were significantly increased, (copper 2-fold and 1.4-fold, and nickel 10.1-fold and 4.3-fold, respectively). Potassium, which is supplemented in BMM and present at much higher baseline levels in untreated cells, was also increased in both conditions (1.4-fold and 1.3-fold, respectively). In KKL-40-treated cells, a 75% decrease in Mn was observed, and in both KKL-40 and KKL-55-treated cells, molybdenum levels were decreased by 50% and 65%, respectively, albeit not with statistical significance in KKL-40-treated cells. Thus, in congruence with the proteomic response, multi-element analysis revealed a disturbance of metal ion homeostasis after treatment with the *trans*-translation inhibitors.

Carrier ionophores such as calcimycin can form complexes with specific metal ions, and these complexes can cross membranes. Given the well-documented toxicity of excessive copper (28), we investigated whether KKL-40 and KKL-55 are capable of forming complexes with Cu(II) using a calcein-based assay. Calcein is a fluorescent dye known to be quenched by copper ions (29). The assay was used on the assumption that if KKL-40 or KKL-55 bind copper, the compound will compete with calcein for copper binding. Different concentrations of KKL-40 or KKL-55 (0 to 30 µM) were added to 3 µM of calcein:Cu(II) complex (Fig. 1d). Fluorescence signals were normalized to calcein fluorescence in the absence of copper. We observed that 50% fluorescence was restored with 5 µM KKL-40 and 10 µM KKL-55, and 100% fluorescence was observed with 12 µM KKL-40 and 30 µM KKL-55. These results indicate that both KKL-40 and KKL-55 form complexes with Cu(II), with KKL-40 competing more efficiently for Cu(II) binding.

We characterized the KKL:Cu(II) complexes with regard to their stoichiometry, copper affinity, and lipophilicity (Table 1). To independently confirm the formation of these complexes, UV-VIS spectra of KKL-40 and KKL-55 were recorded in both HEPES buffer and BMM with and without equimolar Cu(II) addition (Supplementary File 1 Supplementary Fig. S2). Indeed, absorbance peaks shifted in the presence of copper. Absorbance of the KKL-40:Cu(II) complexes peaked at 295 nm, that of KKL-55:Cu(II) at 270 nm. The stoichiometries of these complexes were determined using the Job’s plot method (30), where different concentrations of Cu(II) were titrated against KKL-40 or KKL-55 to maintain a constant total concentration of the complex components (Supplementary File 1 Supplementary Fig. S3). The Cu mole fractions at the infliction point are indicative of KKL-40 forming mostly 3:1 KKL-40:Cu(II) complexes and KKL-55 forming mostly 2:1 KKL-55:Cu(II) complexes. Association constants for KKL-40 or KKL-55 and Cu(II) were determined by competition equilibrium titrations against the colorimetric indicator 4-(2-pyridazo)-resorcinol (PAR) with a known 1:1 stoichiometry and a reported dissociation constant (K_D_) of 2.6 × 10^-15^ M^-1^ at pH 7.4 (31). PAR was sourced at a five-fold molar excess to copper. The K_A_ values at the equivalence point (100 µM PAR, 20 µM Cu(II), 35.15 µM KKL-40 or 185.51 µM KKL-55) were calculated considering the stoichiometries of the complexes (31). At pH 7.4, the K_A_ determined for KKL-40:Cu(II) was ∼ 1.7 × 10^-3^ M^-3^, that for KKL-55:Cu(II) was ∼ 8.6 × 10^-12^ M^-2^ (see Supplementary File 1 Supplementary Fig. S4 for representative titrations). The infliction points for KKL-40 and KKL-55 at which 50% of copper is no longer bound to PAR, were 35.15 µM and 185.51 µM, respectively, indicating a higher copper affinity for KKL-40 than for KKL-55. This is corroborated by the calcein fluorescence quenching assay (Fig. 1d), showing complete quenching at a 3:1 KKL-40:Cu(II) molar ratio, but only approx. 25% quenching at a 2:1 KKL-55:Cu(II) molar ratio.

**Table 1.** Chemical properties of KKL-40:Cu(II) and KKL-55:Cu(II) complexes. Stoichiometry was investigated using the Job’s plot method (30). Association constants (K_A_) were assessed by titration against 4-(2-pyridylazo)resorcinol (PAR). Lipophilicity was determined based on the distribution coefficients in *n*-octanol/buffer. All experiments were performed in HEPES buffer at pH 7.4. Data represent means and SD of three technical replicates.

| KKL | stoichiometry<br>KKL:Cu<sup>II</sup> | association constant<br>[Cu<sup>II</sup>L] $K_A$ | lipophilicity | |
| --- | --- | --- | --- | --- |
| | | | [KKL] $\log D (K_{ow})$ | [Cu<sup>II</sup>L] $\log D (K_{ow})$ |
| KKL-40 | 3:1 | $1.71 \times 10^{-3} \text{ M}^{-3}$ | $0.3 \pm 0.003$ | $1.18 \pm 0.02$ |
| KKL-55 | 2:1 | $8.58 \times 10^{-12} \text{ M}^{-2}$ | $0.74 \pm 0.13$ | $1.58 \pm 0.3$ |

To assess if complex formation might change the ability of KKL-40 and KKL-55 to cross lipid membranes, the lipophilicity was determined for free and complexed molecules. We measured extinction coefficients in *n*-octanol/buffer to determine levels of free and complexed KKL-40 and KKL-55 in the organic and aqueous phases (Supplementary File 1 Supplementary Table S1). Since molecules and complexes may be ionized, the distribution coefficient (*D*) was calculated using equations 2 and 3 (see method section) to describe lipophilicity. Positive Log *D* values indicate lipophilicity and negative values hydrophilicity. Log *D* of KKL-40 increased from 0.3 to 1.18 when complexed with Cu(II) and Log *D* of KKL-55 increased from 0.74 to 1.58 (Table 1). These results indicate that both complexes are more lipophilic than the respective free molecules, increasing the possibility of them interacting with and crossing lipid membranes. Both distribution coefficients are similar to those of the known copper chelating agents 8-hydroxyquinoline and pyrithione (32).

To assess the ability of KKL-40 and KKL-55 to facilitate Cu(II) transport across model membranes, a calcein quenching assay was established using calcein-loaded large unilamellar vesicles (LUV) (Fig. 1e, Supplementary File 1 Supplementary Fig. S5). When lytic concentrations of Triton X-100 (0.2%) were added to the calcein-loaded LUVs, setting free the calcein, no change in fluorescence was observed, indicating that the calcein concentrations used are not self-quenching. However, when 3 µM CuCl_2_ was added simultaneously with Triton X-100, calcein fluorescence was fully quenched due to the calcein released by LUV lysis readily binding Cu(II). In contrast, only a slight decrease in fluorescence was observed when 3 µM CuCl_2_ was added to intact LUVs, indicating that copper enters the LUVs slowly. Importantly, addition of 3 µM CuCl_2_ to LUVs preincubated with 1 µM KKL-40 or 1 µM KKL-55 resulted in rapid quenching of calcein fluorescence, indicating accelerated copper influx facilitated by both compounds. KKL-55 exhibited more pronounced quenching, possibly due to differences in stoichiometry. To exclude the possibility that the quenching was due to LUV lysis, we assessed LUV integrity during exposure to KKL-40 and KKL-55 using vesicles loaded with a self-quenching concentration of calcein (Supplementary File 1 Supplementary Fig. S6). As demonstrated for Triton X-100 treatment, in this case LUV lysis caused an increase in fluorescence due to calcein dilution in the assay volume and the consequent cessation of self-quenching. No increase in fluorescence signal was observed for either KKL-40 or KKL-55, confirming that these molecules do not disrupt LUV integrity. We conclude from the LUV experiments that both KKL-40 and KKL-55 effectively mediate the transport of copper ions across model membranes *in vitro*.

### 2. Co-treatment of *B. subtilis* with CuCl_2_ and KKL-40 or KKL-55 causes synergetic transport and stalls growth

As shown above (in Fig. 1c), an increase in cellular copper levels was observed when *B. subtilis* grown in BMM was treated with KKL-40 or KKL-55 concentrations that inhibit growth of log-phase cells by ∼50%. In contrast to BMM, which contains only trace amounts of copper, normal total copper levels in human blood range from 10.5 to 16.4 µM (33). Given that copper plays a role in the innate antimicrobial immune defense (reviewed in (34)), we explored the effects of copper on the antibacterial effectiveness of KKL-40 and KKL-55. For co-treatments, we chose 3.3 µM CuCl_2_, which by itself is completely benign for *B. subtilis*. When *B. subtilis* was treated simultaneously with 3.3 µM CuCl_2_ and 0.8 µM KKL-40 or 7.2 µM KKL-55 growth stalled immediately (Fig. 2a,b). Growth arrest was also observed when either KKL-40 or KKL-55 was applied at 1 µM in the co-treatment (Supplementary File 1 Supplementary Fig. S7). Since neither 1 µM KKL-55 nor 3.3 µM CuCl_2_ by themselves had any effect on growth, this co-treatment experiment revealed strong synergetic effects.

**Figure 2.**
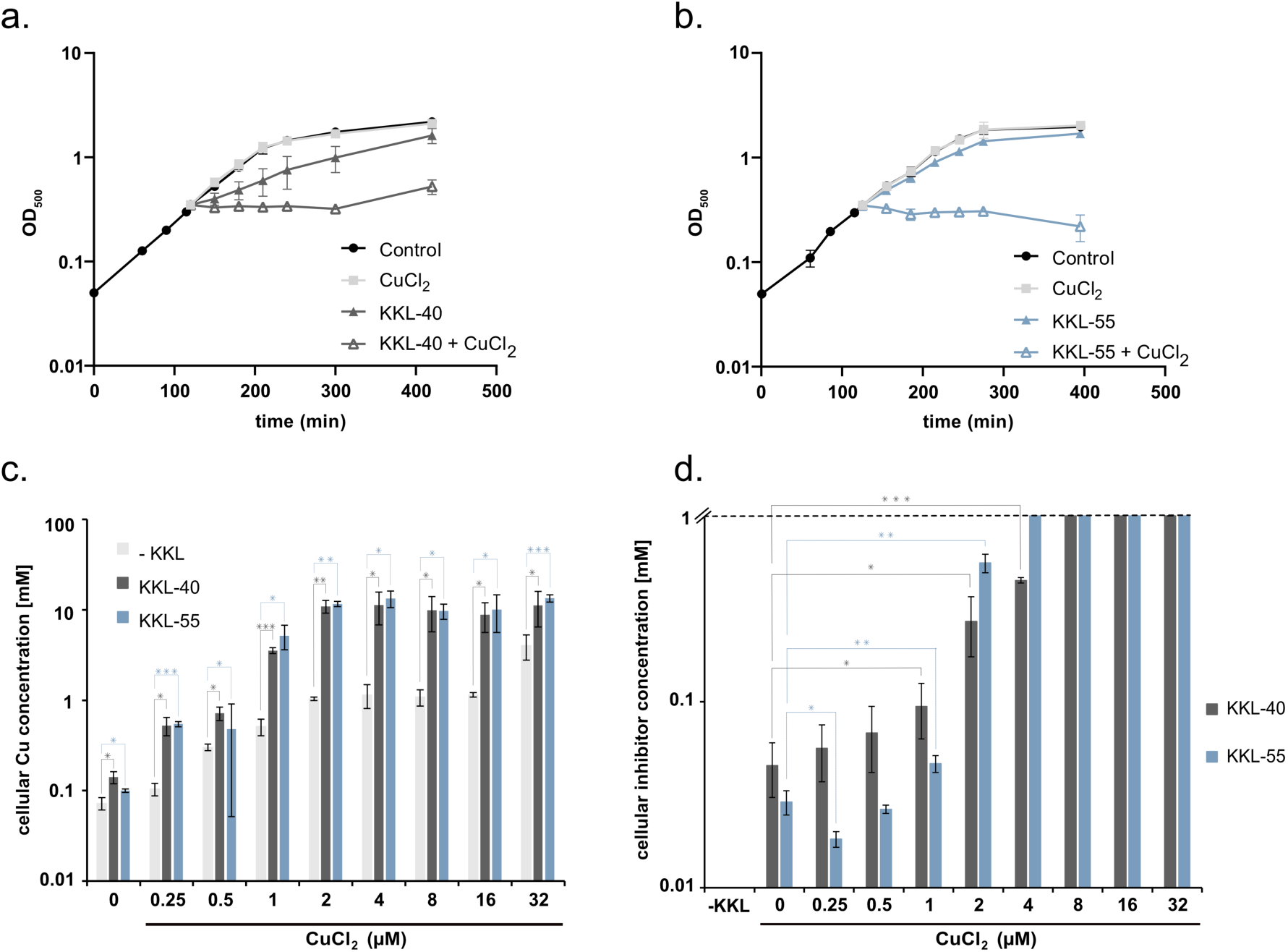
Effect of KKL-40 or KKL-55 and Cu(II) co-treatment on *B. subtilis* growth, copper and KKL levels. (a, b) Cultures were split at an OD_500_ = 0.35 and treated with subinhibitory concentrations of 3.3 µM CuCl_2_ and/or 0.8 µM KKL-40 (a) or 7.2 µM KKL-55 (b). Growth was monitored for three hours after treatment. (c, d) Cellular copper (c) and KKL-40 or KKL-55 (d) concentrations were determined by ICP-OES and mass spectrometry, respectively, 15 min after co-treatment with 0.8 µM KKL-40 or 7.2 µM KKL-55 and different concentrations of CuCl_2_. Cellular KKL-40 and KKL-55 levels exceeded the calibrated range (up to 1 mM) when CuCl_2_ was added at or above 4 µM (see also Supplementary File 2). Data (a-d) represent the means and SD of three biological replicates. Statistical analysis for (c) and (d) was performed using Student’s *t*-test (^✱^ p ≤ 0.1; ^✱✱^ p ≤ 0.01; ^✱✱✱^ p ≤ 0.001).

To further characterize the synergism, we simultaneously treated *B. subtilis* with physiologically effective concentrations of KKL-40 or KKL-55 in combination with copper concentrations ranging from 0.25 µM to 32 µM (Fig. 2c,d). Determination of cellular copper and KKL-40 and KKL-55 levels by ICP-OES and mass spectrometry, respectively, revealed a strong co-dependence of copper and inhibitor transport. After 15 min of treatment with a physiologically effective concentration of KKL-40, the cellular KKL-40 level was 0.045 mM when no CuCl_2_ was added, but ∼0.387 mM when 3.3 µM CuCl_2_ was included with KKL-40. Cellular KKL-55 levels increased even more strongly, from 0.029 mM when no CuCl_2_ was added to >1 mM when 3.3 µM CuCl_2_ was added. This increase in inhibitor levels might be sufficient to explain the growth arrest, since a similarly abrupt and complete growth arrest was observed in the absence of copper supplementation when *B. subtilis* was treated with 10 µM KKL-40 or 36 µM KKL-55 (Supplementary File 1 Supplementary Fig. S8).

We next asked whether the increase in cellular copper is also sufficient to explain the antibacterial effects of KKL-40 and KKL-55. In cells treated with CuCl_2_ and KKL-40 or KKL-55 an increase in cellular copper was observed that exceeded the increase in cells treated with copper only (Fig. 2c, Supplementary File 2). The cellular copper concentration was 0.072 mM in untreated cells, and ∼1.1 mM in cells treated with 3.3 µM CuCl_2_. The concentration increased to ∼11.2 mM in cells treated with KKL-40 and CuCl_2_, and to 12.8 mM in cells treated with KKL-55 and CuCl_2_ (approximate values were estimated by interpolation). However, the high cellular copper levels alone seem not sufficient to explain the growth arrest. In the absence of KKL-40 and KKL-55, the physiologically effective concentration at which CuCl_2_ caused an approximately 50% reduction in growth rate in BMM is 2.98 mM (Supplementary File 1 Supplementary Fig. S9a). At this CuCl_2_ concentration, copper accumulates to even higher levels in the bacteria: 21.5 mM at 15 min and 28.5 mM at 60 min after copper addition (Supplementary File 1 Supplementary Fig. S9b, Supplementary File 2). Since growth was only reduced by about 50% at >20 mM cellular copper, the inhibitor-mediated increase in copper levels only partially explains the antibacterial effects of KKL-40 and KKL-55.

If in the co-treatment the rapidity of copper influx is the main factor contributing to the antibacterial effect, pre-exposure to copper might protect cells from KKL-40 and KKL-55 treatment by triggering the upregulation of the copper stress response resulting in physiological pre-adaptation. If, however, inhibitor influx, rather than copper influx, is the decisive antibacterial factor, one would postulate that prior exposure to copper will not protect *B. subtilis* from the antibacterial effects of KKL-40 or KKL-55 exposure. In a pre-adaptation experiment, 3.3 µM CuCl_2_ was added to *B. subtilis* cultures at different time points prior to the addition of KKL-40 or KKL-55 during early log-phase (Supplementary File 1 Supplementary Fig. S10). As with simultaneous CuCl_2_ and KKL-40 or KKL-55 addition, growth stalled immediately upon inhibitor addition, suggesting that exposure to benign copper concentrations does not prepare the cells to better tolerate inhibitor exposure. This was also observed, when cells were pre-exposed to physiologically effective concentrations of CuCl_2_ (2.98 mM) prior to KKL-40 or KKL-55 treatment (Supplementary File 1 Supplementary Fig. S11).

After addition of 1 µM KKL-40 or KKL-55 (instead of the physiologically effective inhibitor concentrations of 0.8 µM KKL-40 and 7.2 µM KKl-55) a similar co-dependency with copper influx was observed (Supplementary File 1 Supplementary Fig. S7d). However, differences between the two compounds became more apparent. The cellular KKL-40 concentration could differ by more than two orders of magnitude from that of KKL-55. In the absence of CuCl_2_ addition, the cellular KKL-55 concentration was 0.003 mM, whereas the cellular KKL-40 concentration was 0.241 mM. Element analysis suggested that when copper availability is limited, KKL-55 influx relies on trace copper present in the medium, while KKL-40 translocates together with nickel (Supplementary File 1 Supplementary Fig. S7d, Supplementary File 1

Supplementary Fig. S12, Supplementary File 2). When copper was available, a copper concentration-dependent influx of KKL-55 was observed (Supplementary File 1 Supplementary Fig. S7d). At 3.3 µM CuCl_2_ and 1 µM KKL-55 co-treatment, cellular KKL-55 was 0.012 mM. In the same cells, copper accumulated to >20 mM, showing that KKL-55 efficiently acts as a copper ionophore under these conditions. This combination was sufficient to cause growth stalling (Supplementary File 1 Supplementary Fig. S7b). The stronger dependence of KKL-55 on copper for uptake was corroborated using bathocuproinedisulfonic acid (BCS) as a copper chelator (chelates Cu^I^ and Cu^II^) in BMM (Supplementary File 1 Supplementary Fig. S13). While *B. subtilis* growth was inhibited by both KKL-40 and KKL-55 at low BCS concentrations, growth inhibition by KKL-55, but not to the same extent as KKL-40, was abolished at higher BCS concentrations. CuCl_2_ addition restored the effectiveness of KKL-55. From these experiments we conclude that KKL-55 influx depends much more heavily on copper than influx of KKL-40.

### 3. Co-treatment of *B. subtilis* with KKL-40 or KKL-55 and CuCl_2_ stalls protein synthesis

Excess copper can catalyze Haber-Weiss and Fenton-like reactions, causing oxidative stress, and it can displace crucial metals such as iron in proteins, disrupting Fe-S clusters (28, 35). It has further been shown to cause protein precipitation *in vitro* in *E. coli* cell lysates as well as in *vivo*, particularly in anaerobically grown *E. coli* (36, 37) Copper can also affect RNA stability by forming 8-oxo guanosine and by causing mRNA and rRNA strand breaks (38, 39). To gain an understanding how, on a physiological level, the *trans*-translation inhibitors synergize with copper to cause a growth arrest, we performed proteome analyses of cells exposed to single and co-treatments. Initially, following the approach used to investigate the proteomic response to KKL-40 and KKL-55 treatment, we performed a 2D gel-based proteome analysis utilizing radioactive pulse labeling. Pulse-labeling with methionine allows us to evaluate both the overall protein synthesis rate of the cultures in a 5-min time window starting 10 min after treatment initiation, and, after 2D-PAGE, the relative synthesis rates of a large fraction of the cytosolic proteins. We investigated the impact of CuCl_2_ at the non-inhibitory concentration (3.3 µM) and at 2.98 mM CuCl_2_ (the physiologically effective concentration causing a ∼50% growth inhibition) (Supplementary File 1 Supplementary Fig. S14), as well as the impact of KKL-40 and KKL-55 at physiologically effective concentrations in combination with 3.3 µM CuCl_2_. Comparing total protein amounts based on Bradford assay in the soluble cellular fractions of copper-treated and untreated cells, no significant difference was observed, indicating that copper-mediated protein precipitation is but a minor contributor to antibacterial copper toxicity in aerobically grown *B. subtilis*. Neither the 3.3 µM nor the 2.98 mM CuCl_2_ treatment affected the total protein synthesis rates (as determined in the soluble fraction of cell lysates), whereas treatment with the physiologically effective concentrations of KKL-40 (0.8 µM) or KKL-55 (7.2 µM) caused synthesis rates to decrease by 75% and 25%, respectively. When growth is stalled by the co-treatment with either KKL-40 or KKL-55 and 3.3 µM CuCl_2_, protein synthesis rates (but not protein content in the soluble fraction) decreased below 2% (Fig. 3a).

**Figure 3.**
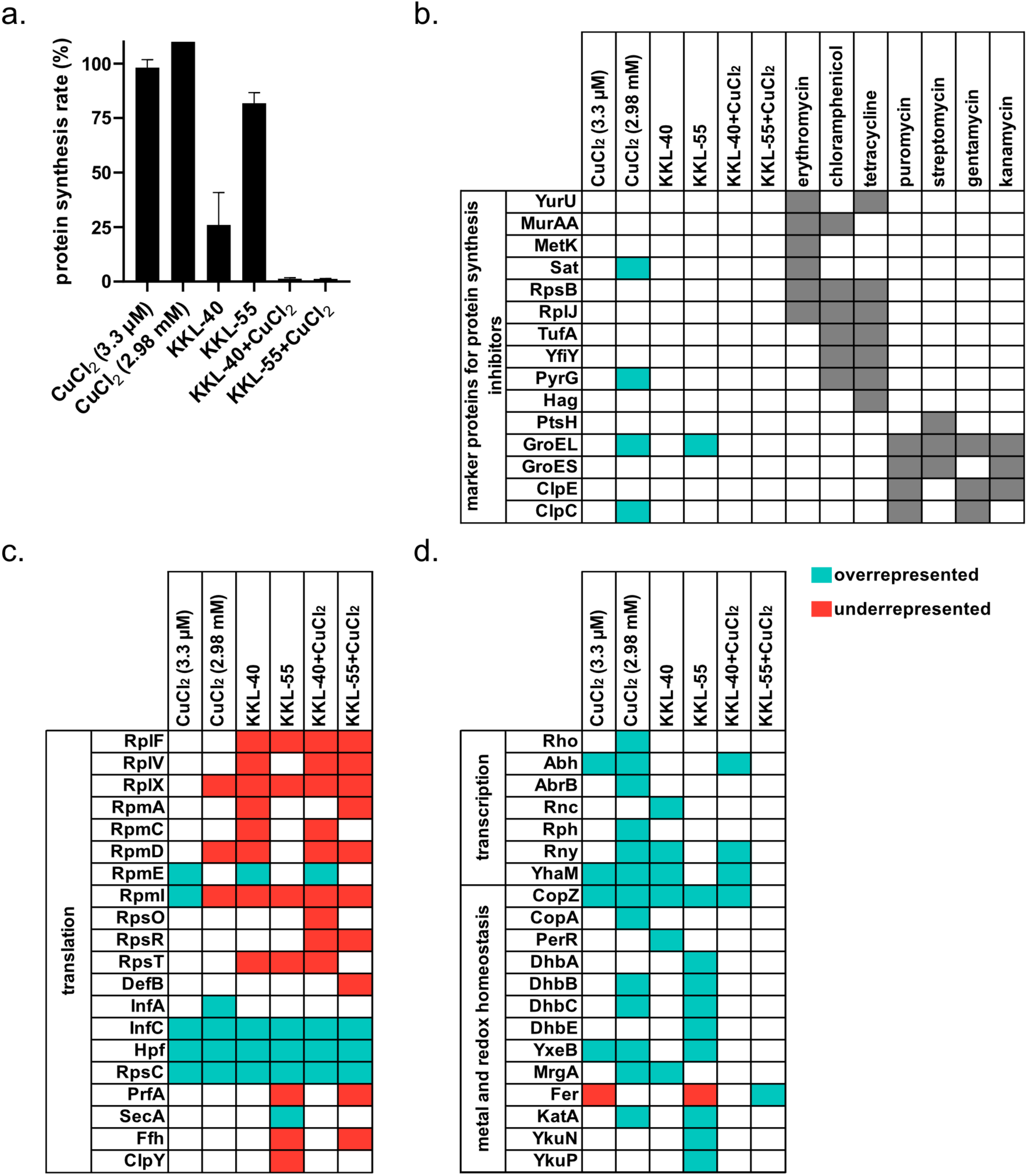
Proteomic response of *B. subtilis* to CuCl_2_, KKL-40, KKL-55, and co-treatments. (a) Protein synthesis rates of cells treated for 15 min with 3.3 µM CuCl_2_, 2.98 mM CuCl_2_, 0.8 µM KKL-40, 7.2 µM KKL-55, 0.8 µM KKL-40 with 3.3 µM CuCl_2_, or 7.2 µM KKL-55 with 3.3 µM CuCl_2_ relative to untreated controls. (b) 2D-PAGE-based comparison of proteomic profiles based on marker proteins identified for known protein synthesis inhibitors. (c,d) Gel-free proteomics. Relative abundance of selected proteins related to protein synthesis (c) and metal and redox homeostasis (d) in samples harvested 60 min after treatment (15 min data in Supplementary File 1 Supplementary Fig. S15, full overview in Supplementary File 3 and Supplementary File 4). Data in (a) represent the means and SD of three biological replicates. In (b,c,d), squares indicate proteins identified as unique or overrepresented (turquoise) or proteins not identified or underrepresented (red) compared to controls. Unique proteins were identified in two or three out of three biological replicates of one condition but not in the other. Overrepresented and underrepresented proteins were selected if their regulation factors exceeded thresholds based on a 95% confidence interval (mean ratio + 1.96 × SD).

As expected, due to the lack of radio-labeled proteins, no proteins were detected on autoradiographs of the 2D-gels for the co-treatments (Supplementary File 1 Supplementary Fig. S14). Treatment with 3.3 µM CuCl_2_, which had no impact on growth, did not elicit any detectable changes in the 2D gel-based proteome profile (Supplementary File 1 Supplementary Fig. S14). In response to 2.98 mM CuCl_2_, however, a number of marker proteins indicative of the cell facing copper stress were upregulated (Supplementary File 1 Supplementary Fig. S14, Supplementary File 1 Supplementary Table S3). Among them were the markers for oxidative stress alkyl hydroperoxide reductase AhpCF, catalase KatA, azoreductases AzoR1 and AzoR2, and nitro/flavin reductase NrfA, as well as general markers for protein stress chaperonin GroEL and protease ClpP. Upregulated were also more specific markers for a disturbance of metal and sulfur homeostasis, including proteins of cysteine, methionine, and sulfur metabolism CysK, SsuD, LuxS, and Sat, the thiamine pyrophosphate-dependent pyruvate dehydrogenase PdhB and thiamin biosynthesis protein ThiC, as well as Zwf (glucose 6-phosphate dehydrogenase) and YugJ, which have Mg^2+^, Mn^2+^, and Ca^2+^ cofactors or a Ni^2+^ cofactor, respectively. The proteomic profile for excess copper conditions was integrated as reference in the tSNE plot (Fig. 1a).

Specifically querying the previously reported proteomic profiles of KKL-40, KKL-55 (13), and the one obtained for 2.98 mM CuCl_2_-treated *B. subtilis* for marker proteins upregulated in response to treatment with known protein biosynthesis inhibitors (Fig. 3b) revealed very little to no overlap that could hint at physiological effects related to the shutdown of translation in co-treatment conditions. In summary, we conclude that the growth arrest in the inhibitor:copper co-treatment conditions goes hand in hand with stalled translation, but we failed to gain insights into potential physiological causes.

We then used a gel-free, label-free quantitative proteomics workflow to assess the proteomes (Fig. 3c,d, Supplementary File 3, Supplementary File 4, Supplementary File 1 Supplementary Fig. S15). This technique provides insights into the relative protein abundance, and a basis for differential analysis even when no translation occurs, and protein abundance is modulated predominantly by degradation and/or aggregation. 60 min after addition of 3.3 µM CuCl_2_ the metallochaperone CopZ was overrepresented (Fig. 3d). Signs of copper stress (28) were observed when cells were treated with 2.98 mM CuCl_2_, including overrepresentation of CopZ and the copper exporter CopA. Beside the oxidative and protein stress response proteins KatA, MrgA and ClpC, proteins involved in the biosynthesis of the siderophore bacillibactin (DhbB, DhbC) accumulated after treatment with CuCl_2_. Bacillibactin production is dependent on iron concentration (40) and up-regulation of the pathway has been observed after copper stress, a reaction attributed to the displacement of iron in Fe-S clusters (41). Furthermore, another previously reported effect of copper toxicity and the related oxidative stress is the formation of mRNA adducts caused by the alkylation and oxidation of the nucleotide bases in the mRNA and RNA cission resulting from reactions with the phosphate sugar backbone (42, 43). These adducts and non-stop mRNAs promote ribosome stalling, requiring ribosome rescue mechanisms for release (44). Here we observed the accumulation of Rny, Rph and YhaM, known polynucleotide phosphorylases or PNPases which can degrade oxidized RNA (45). In congruence with protein synthesis rates being unaffected by the copper treatments, there were few changes in abundance of ribosomal proteins (Fig. 3c). Interestingly, initiation factor 3 (InfC) and hibernation promoting factor (Hpf), and ribosomal protein S3 (RpsC), which localizes at the interface of hibernating ribosomes (46), were at or below the detection limit under control conditions, but overrepresented after copper treatments.

Congruent with the observed decreased protein synthesis rates (Fig. 3a), in *B. subtilis* treated with either KKL-40 or KKL-55 many of the ribosomal proteins were underrepresented, while initiation factor 3 (InfC), hibernation promoting factor (Hpf), and ribosomal protein S3 (RpsC) were overrepresented. Among the ribosomal proteins underrepresented after KKL-40 treatment was bl27 (RpmA), a ribosomal protein whose movement allows for the translocation of tmRNA-SmpB to the P-site of the ribosome (19). Previous studies found that a KKL-40-like acylaminooxidiazole-derived KKL located in non-stop ribosomes stabilizes RpmA in a conformation that inhibits *trans*-translation (19). The destabilization of RpmA observed here, by contrast could be an indication of the degradation of stalled ribosomes. In cells treated for 60 min with KKL-55 release factor 1 (PrfA), which recognizes the UAA/UAG stop codons and catalyzes the release of the nascent protein from the tRNA (47), was no longer detected, likely contributing to limiting translation, while PrfA levels were unchanged after treatment with KKL-40. Consistent with the observed increase in cellular copper levels, in both KKL-40 and KKL-55-treated cells the copper chaperone CopZ is overrepresented. In KKL-40-treated cells, which accumulate higher copper levels, however, the global metal and redox homeostasis transcriptional regulator PerR was overrepresented together with mini-ferritin MrgA, and the ribonucleases Rnc, Rny, and YhaM indicative of copper-related RNA stress (see above), while in KKL-55-treated cells a different set of indicators of disturbed metal ion and redox homeostasis were overrepresented including siderophore production (DhbABCE) and hydroxamate siderophore transport (YxeB) proteins and catalase KatA.

The co-treatment of *B. subtilis* with KKL-40 or KKL-55 and 3.3 µM CuCl_2_ caused protein synthesis to stall, so that most of the observed shifts in protein abundance are likely related to protein stability (and solubility) during co-treatment. As for the single treatments, ribosomal proteins were underrepresented, InfC, Hpf, and RpsC overrepresented, and release factor 1 (PrfA) only detected in co-treatments with KKL-40, but not KKL-55. The copper chaperone CopZ and the ribonucleases Rny and YhaM were overrepresented after KKL-40:CuCl_2_ co-treatment, but in neither co-treatment there was a broad overrepresentation of proteins addressing the disturbance of metal homeostasis and/or oxidative stress that would have required new protein synthesis. Taken together, we conclude from the proteome analysis that under co-treatment conditions ribosomes remain stalled, hibernate or are being degraded causing protein synthesis to stall, thus preventing any dedicated stress response. Given the copper-promoted influx of KKL-40 and KKL-55 (Fig. 2d), this could be a consequence of elevated inhibitor levels under co-treatment conditions.

### 4. Distinct models of synergism are proposed depending on differences in the mechanisms of action of KKL-40 and KKL-55

When undisturbed, *trans*-translation ensures not only the rescue of stalled ribosomes, but also the degradation of faulty mRNAs and incomplete nascent polypeptides (48) (Fig. 4a). Based on the proteome analysis we propose the following models for the synergistic antibacterial action of *trans*-translation inhibitors and copper (Fig. 4b). When at physiologically effective concentrations of KKL-40 *trans*-translation is partially inhibited by KKL-40 binding to the 23S rRNA, the ribosomal A-site is inaccessible both to the ternary complex as well as alternative rescue factors resulting in stalled ribosomes not being rescued efficiently (Fig. 4c). *B. subtilis* countermeasures include the hibernation of ribosomes that are not yet trapped and the elimination of RNAs, both contributing to limiting translation. The upregulation of the repressor PerR effectively represses a metal and oxidative stress response. When physiologically effective concentrations of KKL-55 limit the formation of the ternary complex, *trans*-translation is partially blocked, but alternative rescue factors remain at play (Fig. 4d). While rescue by RqcH leads to the elimination of incomplete nascent polypeptides, faulty mRNAs are not removed (49). BrfA and RF2 cooperate to rescue the ribosome, but both faulty polypeptides and mRNAs are released (8). In contrast to KKL-40 treatment, for KKL-55 treatment the ribosome silencing factor RsfS, which binds to the large ribosomal subunit and prevents subunit joining (50) and prevents 50S dimerization when the alternative RQC system is active (51), is not detected 15 min after treatment initiation (Supplementary File 1, Supplementary Fig. S15) and translation rates at 15 min remain higher, potentially reflecting a delay in ribosome hibernation. The proteome response of KKL-55-treated cells reveals challenges with proteostasis and metal and redox stress response proteins are upregulated, likely triggered by the KKL-55-mediated copper influx.

**Figure 4.**
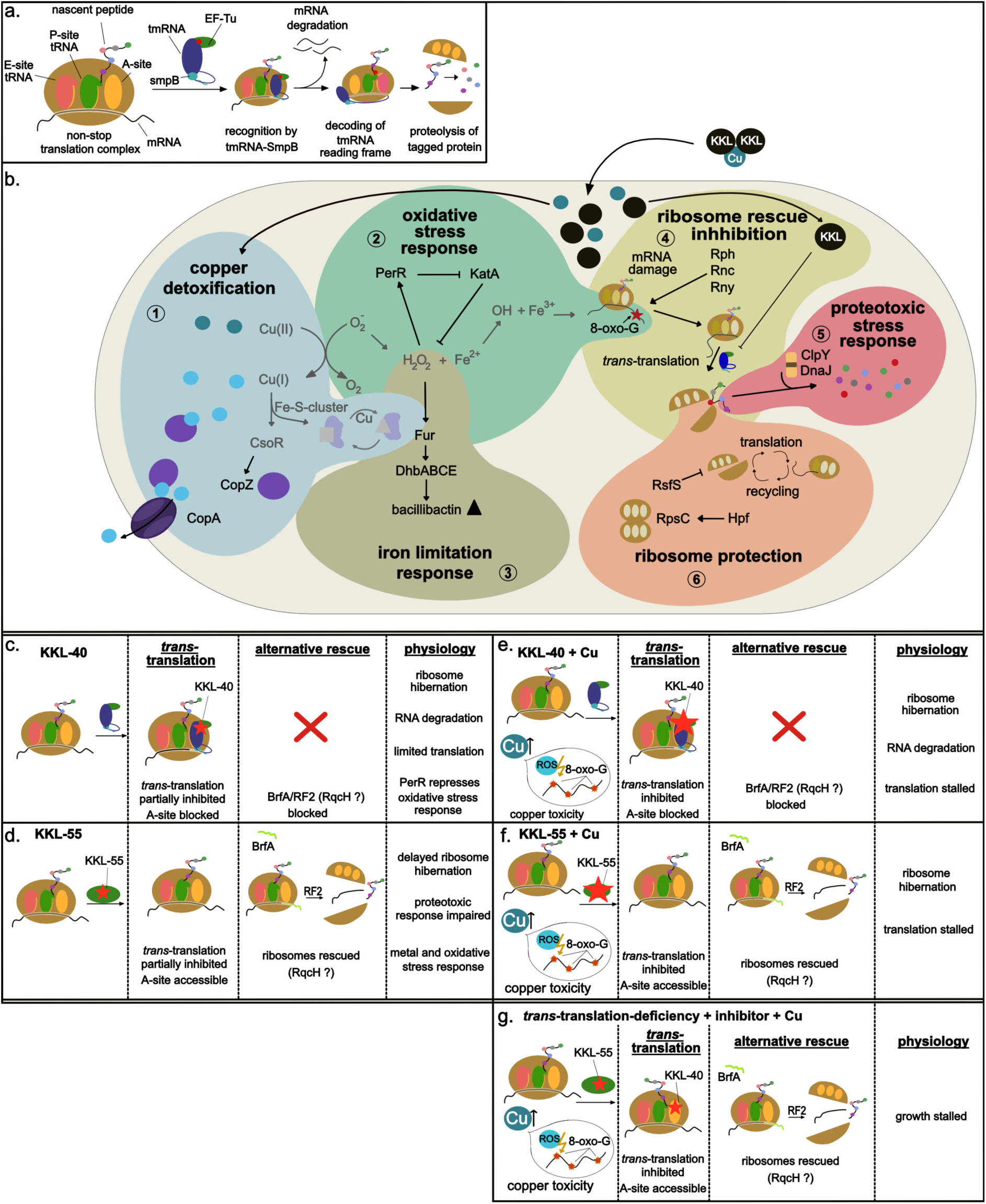
Proposed model for the mode of action of KKL-40 and KKL-55 in *B. subtilis* without and with 3.3 µM CuCl_2_. (a) When *trans*-translation is undisturbed, ribosomes are efficiently rescued, incomplete polypeptides subjected to proteolysis and faulty mRNAs targeted for degradation. (b) KKL-40 and KKL-55 inhibit *trans*-translation and cause a rapid influx of Cu(II). The toxic effects of excess copper include oxidative stress, damage to nucleic acids and proteotoxic stress. Oxidative mRNA damage increases ribosome stalling and the need for ribosome rescue. In treated cells, copper detoxification (**1**) is upregulated, in KKL-40-treated cells the oxidative and metal limitation response (**2** and **3**, respectively) are repressed, while in KKL-55-treated cells both are upregulated. Ribonucleases (**4**) counteracting RNA damage are upregulated in KKL-40, but not KKL-55-treated cells. Proteins protecting from proteotoxic stress (**5**) are down-regulated in KKL-55-treated cells, but not KKL-40-treated cells. Both inhibitors lead to ribosome hibernation (**6**), KKL-40 more efficiently so. (c) Treatment with the physiologically effective concentration of KKL-40 (0.8 µM) causes a partial *trans*-translation block. We hypothesize that the A-site is blocked by the ternary complex that gets trapped due to KKL-40 binding 23S rRNA. Ribosomes hibernate, translation is limited, ribonucleases eliminate faulty RNAs, and PerR represses the oxidative and metal limitation response. A mild copper stress response is observed. (d) Treatment with the physiologically effective concentration of KKL-55 (7.2 µM) causes a partial block of *trans*-translation, but the A-site remains accessible for alternative rescue systems that do not feature degradation of faulty mRNAs and/or incomplete polypeptides. A mild copper stress response as well as oxidative stress and metal limitation responses are upregulated. Some proteins of the proteotoxic stress response are down-regulated and ribosome hibernation is triggered, albeit more slowly than by KKL-40. (e) The co-treatment with KKL-40 and 3.3 µM CuCl_2_ causes a rapid influx of inhibitor and copper. Consequently, *trans*-translation is blocked efficiently. The ternary complex blocks the A-site preventing alternative rescue, translation shuts down due to ribosome stalling and ribosome hibernation, and ribonucleases are stabilized. (f) Co-treatment with KKL-55 and 3.3 µM CuCl_2_ also causes cellular inhibitor and copper concentrations to increase rapidly. Alternative rescue mechanism are active, ribonucleases are not stabilized. Translation stops due to ribosome hibernation and to ribosome rescue capacity being maxed out. (g) *Trans*-Translation-deficient mutants lack the ternary complex so that the A-site remains accessible for alternative rescue when cells are treated with either of the inhibitors and CuCl_2_. The mutants are unable to overcome copper toxicity and growth stalls.

When copper is supplemented at 3.3 µM to the medium, KKL-40 and KKL-55 are taken up much more effectively (Fig. 2d, Supplementary File 1, Supplementary Figure S7d). Uptake appears to be driven primarily by the high number of available cellular copper binding sites in cells. Based on the observed molar increases in KKL-40 and KKL-55 and copper and the proposed complex stoichiometries, KKL-40 and KKL-55 deliver an average of ∼15 and 4400 copper atoms as ionophores before binding their molecular targets. It should be noted, that structural analyses provided no evidence of inhibitor:copper complexes engaging in target interaction. Copper or space for copper ions were neither observed in cryogenic-EM analysis of the interaction of MBX-4132 (structurally related to KKL-40) with the 23S rRNA (19) nor in X-ray analyses of KKL-55:EF-Tu interactions (20). In cells co-treated with KKL-40 and CuCl_2_, levels of cellular KKL-40 free to engage in target inhibition are high enough to very efficiently block ribosome rescue, and RNases are stabilized, leading to a complete shutdown of translation without a significant release of aberrant mRNAs or proteins (Fig. 4e). In the long run, RNases may degrade affected mRNAs and 23S rRNA, contributing to cells recovering and resuming growth (Fig. 2a, Supplementary File 1 Supplementary Fig. S10). Similarly, in KKL-55 and CuCl_2_ co-treated cells *trans*-translation is blocked efficiently by free cellular KKL-55, but ribonucleases are not stabilized leaving mRNAs to be targets of copper damage, and cells are overwhelmed by protein and RNA stress caused by the excess copper and the release of aberrant mRNAs and proteins by alternative rescue systems (Fig. 4f). Translation ceases, due to ribosome hibernation and potentially the trapping of ribosomes in polysomes on faulty mRNAs.

*Trans*-Translation-deficient mutants lacking *ssrA* or *smpB* are unable to form the ternary complex needed for *trans*-translation. Using these *trans*-translation-deficient mutants rather than the small-molecule inhibitors that come with the liability of copper co-transport, we addressed the question of whether or not *trans*-translation plays a role in counteracting copper stress. We compared the copper tolerance of wild-type *B. subtilis* to that of isogenic *ssrA* and *smpB* deletion mutants by determining the minimal inhibitory concentrations of CuCl_2_ (Supplementary File 1 Supplementary Table S3). The MICs for both mutant strains were 0.75 mM lower than that of the wild type, but overall still high (4.47 mM compared to 5.22 mM). This indicates that, in the absence of ionophoric activity, *trans*-translation plays a rather minor role, and that the mutants’ adaptation to slower ribosome rescue (52) still affords growth at comparably high copper concentrations.

We then investigated if *trans*-translation-deficient *B. subtilis* lacking *ssrA* or *smpB* are resistant to KKL-40 and KKL-55 treatment. Especially for KKL-55 we expected it might, since KKL-55 targets EF-Tu preventing formation of the ternary complex that does not form in the mutants. However, the minimal inhibitory concentrations revealed that the mutants are equally sensitive to treatment with KKL-40 and KKL-55 (Supplementary File 1 Supplementary Table S3). They were also vulnerable to KKL-40 or KKL-55 and CuCl_2_ co-treatment during mid-log phase (Supplementary File 1 Supplementary Fig. S16). The high KKL-55 sensitivity is consistent with the proposed model for synergism with copper. KKL-55 acting as ionophore, causes a quick-onset copper toxicity in a system already suffering RNA and protein stress, thereby overwhelming stress response capacities. The high sensitivity to KKL-40 is consistent with observations made for *L. pneumophila*, for which a higher sensitivity of *ssrA* and *smpB* deletion mutants to the structurally related KKL-35 was reported (21). It is also compatible with the proposed model of synergy, if we assume that in the mutants - due to the absence of the ternary complex - binding of KKL-40 to the 23S rRNA no longer prevents alternative rescue factors from interacting with the ribosomal A-site. With only the alternative rescue factors at play, the mutants become as sensitive to the KKL-40-mediated rapid copper influx as they are to KKL-55-mediated influx (Fig. 4g).

Taken together, the loss of *trans*-translation functionality does not seem to be a viable resistance mechanism against copper-ionophoric inhibitors of *trans*-translation. In fact, in conjunction with a copper-ionophoric activity, the specific blocking of *trans*-translation appears preferable to broadly blocking all ribosome rescue. When *B. subtilis* was treated with KKL-40 or KKL-55 in mid-log phase, and then copper was added in addition at different timepoints after the addition of the inhibitor, cells pre-treated with KKL-55 still suffered a complete and lasting growth arrest (Supplementary File 1 Supplementary Fig. S10). This was not the case for KKL-40. The later CuCl_2_ was added, the faster the growth arrest was overcome. At this time, we can only speculate as to why. A possible explanation might be that translation was already stalled effectively enough to reduce vulnerability to the detrimental effects of an additional influx of ribosome rescue inhibitor and copper. An alternative explanation could be that due to the high conditional copper binding affinity of KKL-40, copper competes with the molecular target (the 23S rRNA) for KKL-40 binding, limiting on-target activity.

### 5. Implications of synergistic antibacterial mechanisms of *trans*-translation inhibitors and transition metals

*Trans*-Translation is a promising target for antimicrobial development considering its presence in many pathogens and its relevance for survival and virulence. Both of the *trans*-translation inhibitors tested here, the oxadiazole KKL-40 and the tetrazole KKL-55, synergize with Cu(II) ions for influx and antibacterial activity. It is known that metals can influence the effectiveness of antibiotics, by either enhancing or preventing their activity (53). Copper for example enhances the antifungal activity of the first-line antifungal fluconazole, which has been shown to chelate copper (54). In a recent study, a triazole derivative of fluconazole was developed that acts as Cu(II) ionophore in *Candida* spp (55). Oxadiazole metal complexes have also been developed for different therapeutic applications including antifungals (56). Looking beyond microbes, saccharinate-tetrazoles that also strongly bind Cu(II) showed increased toxicity against tumor cells that was attributed to copper sequestration (57).

The synergy between copper influx and *trans*-translation inhibition appears particularly intriguing, because it is grounded in synergetic mechanisms of action going far beyond a reciprocal potentiation of uptake. The RNA damage caused by the excess copper increases the need for ribosome rescue, and alternative rescue systems – if they kick in – further add to the stress by releasing faulty mRNAs and proteins.

Generally, antibiotics with dual mechanisms are of great interest since resistance and tolerance development in bacteria is slower (58). To the best of our knowledge, no resistance to the KKLs has been observed to date. The sensitivity of *trans*-translation-deficient mutants to copper, and even more so to KKL-40 and KKL-55 treatment, indicates that a *trans*-translation loss-of-function mutation is not a route to resistance despite the different molecular targets.

The synergetic action might also be promising regarding the further development of azol-based *trans*-translation inhibitors. Our growth experiments showed that treatment with KKL-40 or KKL-55 together with an otherwise completely benign concentration of CuCl_2_ (3.3 µM) immediately stalled growth of *B. subtilis*. Normal copper levels in human blood range from 10.5 to 16.4 µM (33) exceeding the supplemented copper concentration. Copper has long been known to play a role in innate immunity (35) and elevated copper levels have been observed e.g. in urine during urinary tract infections (59). Such synergistic action between copper and azol-based *trans*-translation inhibitors might therefore be expected in the context of an infection. Potentially copper supplementation could further enhance the synergetic inhibitory activity as previously proposed for other antimicrobial agents such as 8-hydroxyquinoline and dithiocarbamates (60, 61). It remains to be investigated if the observed synergies extend to pathogens, some of which do not tolerate such high copper concentrations as the soil bacterium *B. subtilis*. *Legionella pneumophila* for example is effectively eliminated within 2.5 h of exposure to 1.57 mM copper solution (62), while *B. subtilis* grows in the presence of 3 mM CuCl_2_ (albeit at lower growth rate).

## Conclusion

Our study indicates a secondary mechanism of action of the *trans*-translation inhibitors KKL-40 and KKL-55. Both are capable of transporting copper across membranes, causing the accumulation of copper in *B. subtilis*. The rapid influx of copper causes protein stress and, more importantly mRNA stress increasing the need for ribosome rescue. The rapid influx of *trans*-translation inhibitors blocks ribosome rescue, terminating translation and stress responses. By importing more copper and allowing alternative rescue, KKL-55 overwhelms the stress response capacities of the cell, resulting in a well-sustained growth arrest. Beyond copper, metal homeostasis is affected more broadly leaving us speculating that additional antibacterial mechanisms remain to be revealed.

## Material and Methods

### Strains, media and antibiotics

*B. subtilis* 168 (DSM 402) (63) was cultivated in chemically-defined Belitzky minimal medium (BMM) (27) containing ultrapure water (ELGA LabWater, UK) with 15 mM (NH_4_)_2_SO_4_ (Sigma Aldrich, USA (purity: ≥ 99%)), 8 mM MgSO_4_ (Grüssing, Germany (purity 99.5%)), 27 mM KCl (Carbolution, Germany (purity: 99%)), 7 mM sodium citrate (Thermo Scientific, USA (purity: 99%)), 50 mM Tris (Roth, Germany (purity: ≥ 99.3%)), 0.6 mM KH_2_PO_4_ (VWR Chemicals, USA (purity: 99.9%)), 2 mM CaCl_2_ (VWR Chemicals (purity: 100%)), 0.001 mM FeSO_4_ (Roth (purity: ≥ 99%)), 0.01 mM MnSO_4_ (Thermo Scientific (purity: 99 %)), 4.5 mM L-glutamate (Sigma Aldrich (purity: 98%)), 0.78 mM L-tryptophan (Carbolution (purity: 98%)), and 11 mM *D*-glucose (Fisher Chemical, USA (purity: 99.5%)), pH 7.5. If not stated otherwise, cultures were grown in flasks at 37°C under steady agitation (200 rpm). Three biological replicates or technical replicates were performed for all experiments as indicated. For biological experiments, KKL-40 (12) and KKL-55 (12) were dissolved in DMSO to a final concentration of 10 mg/ml.

### 2D-PAGE-based analysis of the proteomic response

*Bacillus subtilis* 168 was cultured in BMM and treated at OD_500_ = 0.35 with either 400 µg/ml (2.98 mM) CuCl_2_ or 0.45 µg/ml (3.3 µM) CuCl_2_ (single treatments), or 0.3 µg/ml (0.8 µM) KKL-40 or 2 µg/ml (7.2 µM) KKL-55 with 0.45 µg/ml (3.3 µM) CuCl_2_ (co-treatments). 2D-PAGE-based proteomic analyses were performed as described previously (64). Briefly, for radioactive pulse-labeling with L-[^35^S]methionine, 5 ml of the bacterial culture were treated with a subinhibitory concentration of CuCl_2_ or a combination of either KKL-40 or KKL-55 and CuCl_2_ for 10 min. Cells were incubated with 0.37 MBq radioactive L-methionine (Hartmann Analytic, Germany) for 5 min, and protein biosynthesis was stopped by adding 100 µg/ml chloramphenicol, 1 mM non-radioactive L-methionine in 10 mM Tris-HCl, pH 7.5, and by transferring the cultures to ice. Cells were harvested, washed with 100 mM Tris-HCl/1 mM EDTA pH 7.5) and sonicated using a VialTweeter (Hielscher, Germany). Cell debris was removed by centrifugation, and protein concentrations were estimated by Bradford assay (59) using Roti NanoQuant (Roth, Germany). To determine incorporation rates of radioactive methionine, proteins were precipitated with 20% trichloroacetic acid, and radioactivity was measured using a scintillation counter (Tri-Carb 2800TR, PerkinElmer). For radioactive gels, 50 µg of protein (300 µg for preparative gels) were loaded onto 24 cm immobilized pH gradient strips, pH 4 to 7 (GE Healthcare, United Kingdom), and left for passive rehydration for 16 h. Proteins were separated by isoelectric focusing in the first dimension using a Multiphor II electrophoresis system (GE Healthcare). In the second dimension, proteins were separated according to molecular weight by SDS-PAGE using the Ettan DaltTwelve system (GE Healthcare). Radioactive gels were dried on Whatman paper and exposed to storage phosphor screens (GE Healthcare). Screens were scanned using a Typhoon Trio^+^ instrument (GE Healthcare) at 633 nm excitation wavelength using a 390 nm emission filter. Non-radioactive gels were stained with ruthenium(II)-tris (4,7) diphenyl-1,10-phenanthroline disulfonate and scanned on the Typhoon Trio^+^ instrument exciting at 532 nm and applying a 610 nm emission filter. Image analysis was performed as described previously by Bandow et al. (65) using Decodon Delta 2D 4.2.1 (Decodon, Greifswald, Germany). Signal intensities of protein spots were normalized to the total signal on the autoradiograph. Relative synthesis rates compared to untreated controls were obtained for individual protein spots. With induction factors at or above 2 in each biological replicate, protein spots were designated “markers” and targeted for identification by mass spectrometry. Proteins from 2.98 mM CuCl_2_ treatment were identified from preparative 2D gels after tryptic in-gel digestion. The marker proteins were determined by an LTQ Velos Pro mass spectrometer equipped with a Nano ESI source (Thermo Scientific). The samples were loaded onto a trap column (Acclaim PepMap RSLC 75 μm × 50 cm, Nano Viper) and eluted with a 0.1% formic acid/acetonitrile gradient (400 μl/min, 5% acetonitrile in 4 min, 5-60% acetonitrile in 20 min) from an analytical column (Pep Map 100, C18, 5 μm, 0.1 × 20 mm) at 40°C and subjected to mass spectrometry. The spectra were recorded in a mass range of 50 to 1800 m/z with a 0.5 s/scan in positive mode. Analysis of the spectrum was performed by Xcalibur processing and instrument control software (Thermo Scientific). The protein identification data was uploaded to the ProteomeXchange Consortium via the PRIDE (66) partner repository with the dataset identifier PXD055613 (Project Name: Translation inhibitors and copper ions synergize for enhanced antibiotic activity by increasing uptake and causing a translation arrest that prevents an upregulation of stress responses, Project accession: PXD055613, Reviewer account details: Username, Password: y8GppU3gV155). The proteomic responses to KKL-40 and KKL-55 obtained by 2D-PAGE were taken from Senges et al. (13) and included in the analyses.

### Comparison of proteomic responses (CoPR)

The comparison of proteomic responses was performed as described previously (13) using the proteomic response library of *B. subtilis* to different agents. Proteomic profiles were taken from previous studies (13, 22). Briefly, regulation factors were derived from radioactive 2D-PAGE analysis for proteins identified by mass spectrometry. Regulation factors were logarithmized, and the profile was compared to the reference library using cosine similarity calculations (for source code see https://github.com/CHRSenges/CoPR). t-Distributed stochastic neighbor (27) embedding was performed using the similarity matrix from comparing proteomic responses. To this end, the t-SNE function from scikit-learn (v. 0.23.2) in python (v. 3.8) was used with a perplexity of 7, a learning rate of 10,000 and 10,000 iterations. The library included the responses to 5% DMSO, 4-Br-A23187, acyldepsipeptide, adriamycin, allicin, amidochelocardin, AN3334, anhydrotetracycline, As_2_O_3_, As_2_O_5_, auranofin, aurein, BA234, bacitracin, Bay 20-2369, calcimycin, CCCP, cerulenin, chelocardin, chloramphenicol, cinnamaldehyde, ciprofloxacin, closthioamid, CuCl_2,_ cWFW, cXRX, daptomycin, daunomycin, diamide, erythromycin, FcPNA, gallidermin, GE2270A, gentamicin, gramicidin S, ionomycin, kanamycin, kirromycin, kirrothricin C, KKL-35, KKL-40, KKL-55, linezolid, mersacidin, monensin, MP159, MP196, MP276, MP66, mupirocin, NAI-107, nalidixic acid, nisin, nitrofurantoin, nitroquinolinoxide, nocathiacin I, norvaline, novobiocin, NV503, platencin, platensimycin, puromycin, RcPNA, rhodomyrtone, rifampicin, salvarsan, squalamine, streptomycin, telithromycin, tetracycline, triclosan, trimethoprim, Triton X-100, tunicamycin, UC41, valinomycin, vancomycin, xanthoangelol, amidochelocardin, anhydrotetracycline. Proteomic responses were taken from Senges *et al*. (13, 22).

### Element analysis

*B. subtilis* 168 subcultures (50 ml) were grown in BMM to early logarithmic phase (OD_500_ = 0.35) and treated with either 0.8 µM KKL-40 or 7.2 µM KKL-55. Untreated cells served as a control. Samples of the soluble cell fraction containing the cytosolic and membrane fraction were prepared (67) and analyzed (68) as previously described to determine the cellular element levels. In brief, cells were harvested by centrifugation (3,000 x g for 2 min), washed twice with 100 mM Tris-HCl/1 mM EDTA (pH 7.5), and resuspended in 10 mM Tris-HCl, pH 7.5. Cells were disrupted by sonication with a VialTweeter (Hielscher). Cell debris containing the cell wall was removed by centrifugation (16,000 x g for 10 min), and the supernatant was dried at 50°C with a vacuum centrifuge (EZ-2, Genevac, UK). Dried samples were dissolved in 2.5 ml 65% nitric acid (Bernd Kraft, Germany) by heating at 80°C for 16 h and diluted 1:4 with ultrapure water. Elemental concentrations of P, S, K, Mg, Na, Ca, Mn, Fe, Cu, Mo, Ni and Zn, were determined by inductively coupled plasma - optical emission spectrometry (ICP-OES) using an iCAP 6500 Duo View ICP Spectrometer (Thermo Fisher Scientific) and calibration standards for each element (Bernd Kraft).

### Calcein-based copper binding assay

Calcein assays were performed in 96-well plates. Calcein (Merck, Germany) was dissolved in 50 mM HEPES, pH 7.4, to a final concentration of 3 µM. CuCl_2_ was added to a final concentration of 3 µM. After 5 min incubation, KKL-40 or KKL-55 were added to reach final concentrations of 0.75, 1.5, 3, 6, 12, 15, and 30 µM. Fluorescence was measured through the bottom of a black clear-bottom microplate at λ_ex_ = 480 nm (λ_em_ = 520 nm) using a EnSpire 2300 plate reader (PerkinElmer, USA). Fluorescence was normalized to calcein fluorescence in the absence of Cu(II). Three technical replicates were performed.

### Cu(II) transport into large unilamellar vesicles

Transport of Cu(II) into vesicles was measured as previously described (22, 69). Briefly, Calcein-filled large unilamellar vesicles (LUVs) were prepared using 5 mg of 1,2-dioleoyl-*sn*-glycerophosphatidylcholine (DOPC; Avanti Polar Lipids, Alabaster, AL, USA) in chloroform, which was dried by rotary evaporation overnight at 25 kPA to produce a lipid film on the wall of a 25 x 150 mm glass tube. The lipid film was hydrated with 1 ml of buffer A (10 mM HEPES pH 7.4, 150 mM KCl) containing 300 µM calcein (Merck), by vortexing in the presence of a 3 mm glass pearl. The resulting suspensions were subjected to five freeze-thaw cycles, followed by 15 cycles of extrusion through 0.2 µm nucleopore polycarbonate membranes (Cytiva Global Life Sciences Solutions, USA) mounted in a mini-extruder (Avanti Polar Lipids). Calcein-filled LUVs were separated from free calcein by size-exclusion chromatography (Sephadex G-50) and stored at 10°C. For Cu(II) transport measurements, 20 µl of purified calcein-filled LUVs and 1 µl of a 20 µM KKL-40 or KKL-55 solution (dissolved in DMSO), or 1 µl of DMSO or 20 µl of Triton X-100 as control, were added to 2 ml of buffer A in a cuvette and incubated for 5 min at 4°C. The final concentration of KKL-40 or KKL-55 was 1 µM and of Triton X-100 0.2% v/v. Fluorescence was measured at λ_ex_ = 480 nm (λ_em_ = 520 nm) at 10°C using a PTI-Quantamaster 800 fluorometer (Horiba, Germany), with 3 nm slit width and 1 s resolution. After 1 min of equilibration, 3 µl of a 2 mM CuCl_2_ solution were added to give a final concentration of 3 µM. Fluorescence was recorded for a total of 10 min. The signal for the released dye was normalized to the highest fluorescence intensity measured in the absence of CuCl_2_. Three technical replicates were performed.

To evaluate the integrity of the vesicles in the presence of KKL-40 or KKL-55, calcein-filled LUVs with self-quenching calcein concentrations (750 µM) were used. Triton X-100 (0.2% v/v) served as a positive control for lysis, causing an immediate increase in fluorescence due to calcein dilution into the assay volume.

### Stoichiometry determination by Job’s plot method

To define the characteristic absorbances of KKL-Cu(II) complexes, KKL-40 and KKL-55 absorbances were monitored in the presence and absence of Cu(II) in 50 mM HEPES buffer (pH 7.4) or BMM (pH 7.5) as indicated, using quartz cuvettes with 1 cm pathlength and a V-750 spectrophotometer (Jasco, Japan). Characteristic wavelengths were 295 nm for KKL-40, 350 nm for KKL-40:Cu(II), 270 nm for KKL-55 and 315 nm for KKL-55:Cu(II). The stoichiometries of KKL-40 and KKL-55 for Cu(II) were assessed by Job’s method of continuous variation. Dilutions containing KKL and Cu(II) were prepared in 50 mM HEPES buffer, pH 7.4, such that [(Cu(II)]+[KKL] remained constant as the mole fraction of Cu(II) varied from 0.1 to 1. UV-Visible absorption was obtained using UV-VIS spectroscopy (Microplate Spectrophotometer Epoch BioTek). Stoichiometry was determined by plotting the absorbance at the characteristic wavelength for each complex with low initial absorbance from the KKL-40 or KKL-55 against the mole ratio of Cu^2+^ ([Cu^2+^]/([Cu^2+^]+[KKL])).

### Determination of Cu(II) conditional association constants

To determine the association constants for Cu(II), the indicator 4-(2-pyridylazo)resorcinol (PAR) (Sigma Aldrich, Germany) was used. PAR forms a 1:1 complex with Cu(II) (*Α*_500_ ε = 37000 M^-1^cm^-1^) with a conditional dissociation constant for Cu(II) at pH 7.4 of 2.6 × 10^-15^ M^-1^ (31). A solution of 100 µM PAR and 20 µM Cu(II) was prepared using CuCl_2_, into which KKL-40 and KKL-55 were titrated. After each addition of 1 µl of 2 mM KKL-40 or KKL-55, the solution was left to equilibrate until no spectral changes were observed. Spectra were acquired using a 1 cm pathlength quartz cuvette and a V-750 spectrophotometer (Jasco). The decrease in absorbance at 500 nm upon addition of inhibitor indicated the dissociation of the PAR-Cu complex and the association of the KKL-Cu complex as change in equilibrium described in equation 1. Average equivalence points were determined based on three technical replicate experiments using GraphPad Prism 10 (version 10.1.1). The KKL and PAR concentrations at the equivalence point (50% dissociation of PAR-Cu) were used to calculate the association constants for KKL-40_3_:Cu and KKL-55_2_:Cu complexes. The calculated binding affinities are the averages and SD of three technical replicates for each inhibitor.

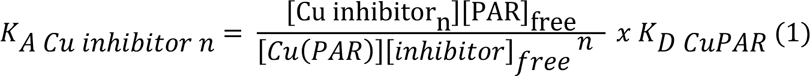

where n = number of inhibitor molecules in the complex, and [Cu inhibitor_n_] = [CuPAR] at the equivalence point (31).

### Partition coefficient

For free inhibitors and inhibitor-Cu(II) complexes, 200 µM solutions were prepared in either *n*-octanol (Sigma Aldrich) or 0.1 M HEPES buffer (pH 7.4). Initial concentrations were determined by UV-VIS absorption spectroscopy using the extinction coefficients in Supplementary Table S1. 4 ml aliquots of each test solution were mixed in a 15 ml Falcon tube with 4 ml of the opposing phase, either *n*-octanol or 0.1 M HEPES (pH 7.4). The mixtures were mixed for 3 h in a 360° rotator. The tubes were centrifuged to separate the phases, and 1 ml aliquots of each phase were removed to measure the UV-VIS spectra (1 cm pathlength quartz cuvettes, V-750 spectrophotometer (Jasco)) from which KKL concentrations were calculated. Assuming free inhibitor to be uncharged and inhibitor-Cu(II) complexes to be charged, the distribution coefficients were calculated according to equations 2 and 3, respectively.

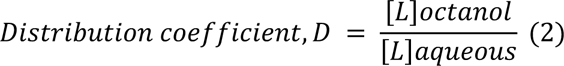

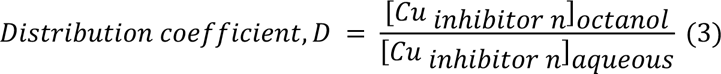

where *n* = number of inhibitor molecules in the complex.

### Growth curve of *B. subtilis* treated with KKL-40 or KKL-55 and CuCl_2_

*B. subtilis* 168 cultures were inoculated to an OD_500_ = 0.05 and left to grow to an OD_500_ = 0.35. Cultures were split and treated with 0.8 µM KKL-40 or 7.2 µM KKL-55 or 1 µM KKL-40 or KKL-55 and 3.3 µM CuCl_2_. Untreated cultures served as controls, cultures exposed only to inhibitor or CuCl_2_ were used for comparison. Cultures were monitored photospectrometrically for three hours after compound addition. Three biological replicate experiments were performed.

### Cellular ion concentrations in cells treated with KKL-40 or KKL-55 and CuCl_2_

*B. subtilis* 168 subcultures were grown in BMM to early logarithmic phase (OD_500_ = 0.35) and treated with 0.8 µM KKL-40 or 7.2 µM KKL-55 or 1 µM of either inhibitor. CuCl_2_ was added to untreated and KKL-treated samples. For 0.8 µM KKL-40 or 7.2 µM KKL-55, final concentrations of CuCl_2_ were 0.25, 0.5, 1, 2, 4, 8, 16, and 32 µM. For 1 µM inhibitor, final CuCl_2_ concentrations were 0.25, 0.5, 0.75, 1, 3, 6, 9 and 18 µM. Untreated cells served as controls. Samples were prepared and analyzed as described above for element analysis. Each sample consisted of 50 ml of culture, and the analysis was conducted in biological triplicates. To calculate cellular concentrations, data was normalized to cell density (OD_500_ = 1 equals 6×10^7^ cells/ml) and a cytoplasmic cell volume of 3.09×10^-9^ µl determined by cryo-electron microscopy by Matias and Beveridge (70, 71).

### Cellular KKL-40 and KKL-55 accumulation in the presence of different Cu(II) concentrations

To determine the cellular inhibitor levels, the cytosolic and membrane fraction was collected and non-covalent target interaction of KKL-40 and KKL-55 was broken up by methanol precipitation. *B. subtilis* 168 subcultures were grown in BMM to the early logarithmic phase (OD_500_ = 0.35). Samples were taken from the same cultures as in the previous sections. 10 ml of cells were centrifuged and washed twice with 100 mM Tris-HCl/1 mM EDTA (pH 7.5) and resuspended in 400 µl methanol. Cells were disrupted by sonication with a VialTweeter (Hielscher). Cell debris was removed by centrifugation after the samples were stored at −20°C for 16 h. 5 µl of supernatant were injected into an ACQUITY UPLC-I-Class system (Waters, USA) equipped with an ACQUITY PREMIER HSS T3 column (particle size 1.8 µm, column dimensions: 2.1 x 100 mm). A gradient with H_2_O and acetonitrile, each with 0.1% formic acid, was used with a flow rate of 0.6 ml/min (Table 2).

**Table 2.** Liquid chromatography gradient. A gradient with H_2_O and acetonitrile (ACN), each with 0.1% formic acid (FA) was used with a flow rate of 0.6 ml/min

| time (min) | % ACN + 0.1 %<br>FA |
| --- | --- |
| 0 | 98.0 |
| 2 | 98.0 |
| 9 | 0.5 |
| 11 | 0.5 |
| 13 | 98.0 |
| 14 | 98.0 |

Data-independent MS^E^ measurements were performed with a Vion IMS QToF (Waters) with an ESI source in positive sensitivity mode. Masses from 50 to 2000 m/z were detected with 0.1 s per scan, and leucine enkephalin was injected as a reference mass every 5 min. Used parameters: capillary voltage, 0.8 kV; sample cone voltage, 40 V; source offset voltage, 80 V; cone gas flow, 50 l/h; desolvation gas flow, 1000 l/h; source temperature, 150°C; desolvation temperature, 550°C; collision gas, N_2_; collision low energy, 6 V; collision high energy ramp, 28-60 V.

KKL-40 and KKL-55 concentrations were determined using a calibration curve of a standard with m/z, ± 2 mDa; retention time, ± 30 s; and CCS, ± 2 %.

### Dependence of the antibacterial activity of KKL-40 and KKL-55 on copper availability

96-well microtiter plates were prepared with BMM containing 0, 10, 50, 100, 250, 500, 1000 and 2000 µM of the copper chelator bathocupronedisulfonic acid (BCS; Sigma Aldrich). Two experiments were performed depending on copper supplementation: one had no copper supplementation, the other was supplemented with 3.3 µM of CuCl_2_. 5 × 10^5^ cells/ml of exponentially growing *B. subtilis* 168 were used to inoculate the prepared plates. Minimal inhibitory concentrations of KKL-40 (14.2 µM) or KKL-55 (18.8 µM) were added. Cells were incubated at 37°C for 16 h. Final absorption at OD_500_ was measured using a Microplate Spectrophotometer Epoch (BioTek, USA) as a readout for growth. Three biological replicates were performed and results normalized to the absorbance of the untreated control of the respective biological replicate.

### Proteomic profiling by quantitative gel-free LC-MS

Proteomic profiling was performed as previously described (52). Quantitative gel-free MS was performed using three biological replicates. Briefly, *B. subtilis* 168 was grown in BMM overnight to exponential growth phase. 50 ml cultures inoculated to OD_500_ = 0.05 were grown and treated at OD_500_ = 0.35 with 3.3 µM CuCl_2_, 2.98 mM CuCl_2_, 0.8 µM KKL-40, or 7.2 µM KKL-55 or co-treated with 3.3 µM CuCl_2_ and KKLs (added independently). Cells were harvested after 15 and 60 min of treatment. Cells were washed and resuspended in 50 mM triethylammonium bicarbonate (TEAB) buffer (Sigma-Aldrich). Cells were disrupted by sonication using a VialTweeter (Hielscher) and centrifuged to remove cell debris. Protein concentrations were determined by Bradford assay (72). 50 µl aliquots containing 0.5 µg/ml of protein were prepared. 0.1% RapiGest (Waters, Milford, MA) was added to each sample to enhance enzymatic cleavage. Cysteines were reduced and alkylated by the addition of 2.5 mM Tris (2-carboxyethyl)phosphine (Sigma-Aldrich) for 45 min at 60°C and then 5 mM iodoacetamide (Sigma-Aldrich) for 15 min at 25°C in the dark. Proteins were digested with trypsin (sequencing grade; Promega, USA) with a protease-to-substrate ratio of 1:100. Digestion was complete after incubation for 5 h at 37°C and 350 rpm. 2 µl of trifluoroacetic acid (Biosolve, Valkenswaard, The Netherlands) were added for RapiGest precipitation and samples were centrifuged at 4°C and 14,000 rpm. Before injection, 50 fmol PhosB peptides were added as a Hi3 quantitation standard (Waters) to 0.5 µg of protein for each sample. Samples were injected into an Acquity M-class ultraperformance liquid chromatography (nanoUPLC) system coupled to a Synapt XS mass spectrometer (Waters).

Proteins were identified as described previously (52). Briefly, data were analyzed using ProteinLynxGlobalServer (PLGS) 3.0.3 software (Waters). Mass spectra were processed using the following parameters: chromatographic peak width, automatic; MS TOF resolution, automatic; lock mass for charge, 785.8426 Da/e; lock mass window, 0.25 Da; low-energy threshold, 200 counts; elevated-energy threshold, 50 counts; elution start time, 60 min; elution end time, 300 min. The reference proteome of *B. subtilis* 168 database (Uniprot Proteome ID: UP000001570, downloaded on March 1st, 2022) containing 4263 protein entries, including the sequences for PhosB, trypsin, and keratin, was used for protein identification using the following parameters; peptide tolerance, automatic; fragment tolerance, automatic; minimal fragment ion matches per peptide, 2; minimal fragment ion matches per protein, 5; minimal peptide matches per protein, 2; maximum protein mass, 250 kDa; primary digest reagent, trypsin; secondary digest reagent, none; missed cleavages, 1; fixed modification, carbamidomethyl C; variable modifications, deamidation N and Q, and oxidation M; false positive rate, 4%; calibration protein PhosB standard; calibration protein concentration for Hi3 quantitation, 50 fmol. The false discovery rate was estimated by searching the shuffled, randomized *B. subtilis* database mentioned above.

Statistical analysis was performed with Microsoft Excel 2020. Protein concentrations were normalized to a six-peptide Phosphorylase B standard (Waters) and subsequently, to total protein on column for each sample. For each biological condition, the mean %SD of protein quantities for the three biological replicates was below 30%. Proteins were selected as unique to the treatments if they were detected in at least two of three replicates but not in the untreated control. Proteins reported as unique to the untreated control were selected using equivalent criteria. Differentially abundant proteins in treated *versus* untreated samples were selected based on log_2_ratios applying a 95% confidence interval (mean ratio + 1.96 × SD) as threshold (Supplementary File 3).

Proteins were assigned to their gene ontology (GO) biological process according to Uniprot and Subtiwiki (73). For each protein, the number of identified peptides, their sequence, counts, location, false discovery rate, and sequence coverage can be found in the files uploaded to PRIDE. The mass spectrometry-based proteomics data was uploaded to the ProteomeXchange Consortium via the PRIDE (66) partner repository with the dataset identifier PXD053533 (Project Name: Translation inhibitors and copper ions synergize for enhanced antibiotic activity by increasing uptake and causing a translation arrest that prevents an upregulation of stress responses, Project accession: PXD053533, Reviewer account details: Username, Password: MyaMfajx2vwU).

## Acknowledgements

We thank Petra Düchting of the chair of Molecular Genetics and Physiology of Plants, Ruhr University Bochum, for her technical support with the ICP-OES measurements. HDU was awarded with a scholarship from the Friedrich Ebert Foundation. JEB would like to thank Barrie Wilkinson (John Innes Centre, UK) for hosting her during the sabbatical. Funding and support were provided by a grant from the National Institutes of Health (R01GM121650) and the German Research Foundation (BA 4193/6-1, RTG 2341 “Microbial Substrate Conversion”). JEB gratefully acknowledges funding for the Synapt mass spectrometer from the German Federal State of North Rhine-Westphalia and the European Union, European Regional Development Fund for the Research Infrastructure Center for System-Based Antibiotic Research (CESAR)(EFRE-0200598) and for the VION mass spectrometer from the German Research Foundation and the German State of North Rhine-Westphalia for the mass spectrometer (“Forschungsgroßgeräte” nach Art. 91b GG, INST 213/961-1 FUGG).

## Conflicts of interest

The authors declare no conflicts of interest.

## Notes

**Competing Interest Statement** Authors declare no conflict of interest.

